# Propidium iodide staining underestimates viability of adherent bacterial cells

**DOI:** 10.1101/475145

**Authors:** Merilin Rosenberg, Nuno F. Azevedo, Angela Ivask

**Affiliations:** Laboratory of Environmental Toxicology; National Institute of Chemical Physics and Biophysics; Akadeemia tee 23, 12618 Tallinn, Estonia; Department of Chemistry and Biotechnology; Tallinn University of Technology; Akadeemia tee 15, 12618 Tallinn, Estonia; LEPABE - Laboratory for Process Engineering, Environment, Biotechnology and Energy; Department of Chemical Engineering; Faculty of Engineering; University of Porto; Rua Dr. Roberto Frias, 4200-465 Porto, Portugal

## Abstract

Combining membrane impermeable DNA-binding stain propidium iodide (PI) with membrane-permeable DNA-binding counterstains is a widely used approach for bacterial viability staining. In this paper we show that PI staining of adherent cells in biofilms may significantly underestimate bacterial viability due to the presence of extracellular nucleic acids. We demonstrate that gram-positive *Staphylococcus epidermidis* and gram-negative *Escherichia coli* 24-hour initial biofilms on glass consist of 76 and 96% PI-positive red cells *in situ*, respectively, even though 68% the cells of either species in these aggregates are metabolically active. Furthermore, 82% of *E. coli* and 89% *S. epidermidis* are cultivable after harvesting. Confocal laser scanning microscopy (CLSM) revealed that this false dead layer of red cells is due to a subpopulation of double-stained cells that have green interiors under red coating layer which hints at extracellular DNA (eDNA) being stained outside intact membranes. Therefore, viability staining results of adherent cells should always be validated by an alternative method for estimating viability, preferably by cultivation.

## Introduction

Propidium iodide (PI) is widely used for bacterial viability staining, especially since Boulos et al. (1999) published the method^1^. PI can only cross compromised bacterial membranes and is therefore considered to be an indicator of membrane integrity. It stains DNA and RNA inside of dead cells or the ones with reversibly damaged membranes. For viability staining PI is usually coupled with a universal stain that crosses intact membranes and stains the DNA and RNA of all cells, thereby enabling to obtain total cell counts. One of the most common examples of such co-stain is SYTO 9. During co-staining with PI and SYTO 9, SYTO 9 can enter all cells regardless of their membrane integrity, bind to DNA and RNA and emit green fluorescence while PI can only enter cells with compromised membranes, bind to DNA and RNA and emit a red fluorescent signal. With higher affinity to bind DNA and in sufficient excess to SYTO 9, PI replaces SYTO 9, when both stains are exposed to the same DNA resulting in red fluorescent signal. As a result of coupling of those two DNA-binding and membrane permeability dependent stains red signals from cells are considered as “dead” and green signals as “alive”^1–3^. Although this principle is widely applied and proven to work well for an array of planktonic cultures, it has its limitations i.e. unequal SYTO 9 staining of viable and dead cells, incomplete replacement of SYTO 9 by PI or energy transfer during co-staining ^2,4^. It has also been demonstrated that PI might in some cases provide false dead signals entering viable cells with high membrane potential ^5^, and that the staining result might be dependent on physiological processes other than membrane damage ^6^. PI-based viability staining results do not always correlate with cultivability also due to the viable but not cultivable (VBNC) state of bacterial cells ^7^ or cell clumping ^8^. Despite its above-mentioned draw-backs, PI and SYTO 9 co-staining is also a widely used and suggested method in biofilm research ^8–17^.

Another factor to consider when staining cells with DNA/RNA-binding fluorophores is that nucleic acids are not always only localized inside bacterial cells and surrounded by a membrane. For example, extracellular DNA (eDNA) can be present in planktonic cultures in specific growth phases ^18^. During biofilm formation, eDNA mediates bacterial attachment to surfaces ^19^, and it also plays a major role in mature biofilms. The importance of eDNA in biofilm formation has been proven by the fact that DNase I inhibits biofilm formation or detaches existing biofilm of several gram-positive and gram-negative bacterial species ^20^. For the same reason, DNase is also proposed to be used as an anti-biofilm agent ^21,22^. The presence of DNA from non-viable sources (eDNA and DNA from dead cells) has also introduced the need to use ethidium monoazide (EMA), propidium monoazide (PMA) or endonuclease (DNase I) treatment prior to viability assessment by quantitative polymerase chain reaction (qPCR) ^23–25^. All the above-mentioned treatment agents, EMA, PMA as well as DNase I are intact membrane impermeable DNA-targeting compounds spatially targeting the same DNA as PI, depending on membrane integrity.

To get an overview whether the presence of eDNA in biofilms has been considered as a factor that may interfere with PI-based fluorescent staining, we performed a search in Scopus database for "biofilm" and "propidium iodide" and received 683 results while adding "extracellular DNA" or "eDNA" to the search decreased the number of results to 43 indicating that while PI is used for staining biofilms, possible presence of eDNA is generally not taken into account in this context. In the literature we can find that PI is also used for staining of eDNA ^26,27^, but no clear quantitative proof about PI not being suitable for biofilm viability staining because of the presence of nucleic acids in biofilm ECM. More surprisingly, viability staining based on intact membrane impermeable DNA-binding stains like PI are occasionally used even while specifically studying eDNA ^28^. Nonetheless, from some of the articles, hints of such threat can be found. For example, Gião and Keevil observed that some of *Listeria monocytogenes* biofilms in tap water and most of the old biofilms grown in rich media stained red with PI and SYTO 9 co-staining, but were cultivable and suspected red staining not to be indicative of dead cells but to be caused by eDNA ^29^. From these sources it could be suspected that PI-based viability staining of biofilms, although commonly used, could be critically affected by eDNA and cause underestimation of biofilm viability. To address this possibility, we performed quantitative viability assessment of adherent cells using various staining and culture-based methods.

## Results

A combination of epifluorescence microscopy (EM), flow cytometry (FCM) and confocal laser scanning microscopy (CLSM) performed on propidium iodide (PI) and SYTO 9 stained adherent and harvested bacterial cells in parallel with culture-based methods was used to reveal whether staining of adherent bacteria with PI may underestimate their viability. Initial (24 h) biofilms of gram-negative *E. coli* K-12 wild-type substrain MG1655 and a gram-positive *S. epidermidis* type strain DSM-20044 were used for the experiments. *E. coli* MG1655 is widely used in molecular biology and capable of forming biofilm under both aerobic and anaerobic conditions ^30–34^. *S. epidermidis* strains have well established biofilm forming properties similarly to *Staphylococcus aureus* and have been shown to produce eDNA ^13,35^. The biofilms of these two bacterial strains on glass surfaces were formed in phosphate buffered saline (PBS) to rule out potential effect of osmotic stress on bacterial membranes and possibly consequently on viability staining outcome.

### Viability staining *in situ*

As can be seen on representative images (Fig. 1) and from quantitative data (Fig. 2), after PI+SYTO 9 co-staining, most adherent cells (96.35±5.3% of *E. coli* and 75.69±18.44% of *S. epidermidis* cells) in 24 h biofilm in PBS stained red with PI *in situ* (Fig. 1a, 1b, 2a, 2b) while most (about 99%,) planktonic cells from suspension above the respective biofilms stained green with SYTO 9 on a filter (Supplementary Fig. 1). This could normally be interpreted as simply showing the differences in the physiology of adhered and planktonic cells and different proportion of dead and alive cells indicating better viability of planktonic cells. However, it is harder to explain this difference taking into account that adherent cells demonstrated biofilm-specific aggregation into microcolonies, no toxic agent was added, samples were rinsed before staining and loose dead planktonic cells should have been removed. Also, the proportion of red-stained cells in the initial biofilms was surprisingly high. For example, using the same staining method, Wang *et al* noted only a few dead cells among viable cells on a 24 h *E. coli* biofilm on silicone in PBS ^36^. Starved biofilms incubated in PBS are more commonly used in oral health studies where most of the cells in biofilm tend to stain green similar to Zhu *et al* reporting 76.7% viability of 24 h *Streptococcus mutans* biofilm on glass in phosphate buffer ^9^.

**Figure 1.**
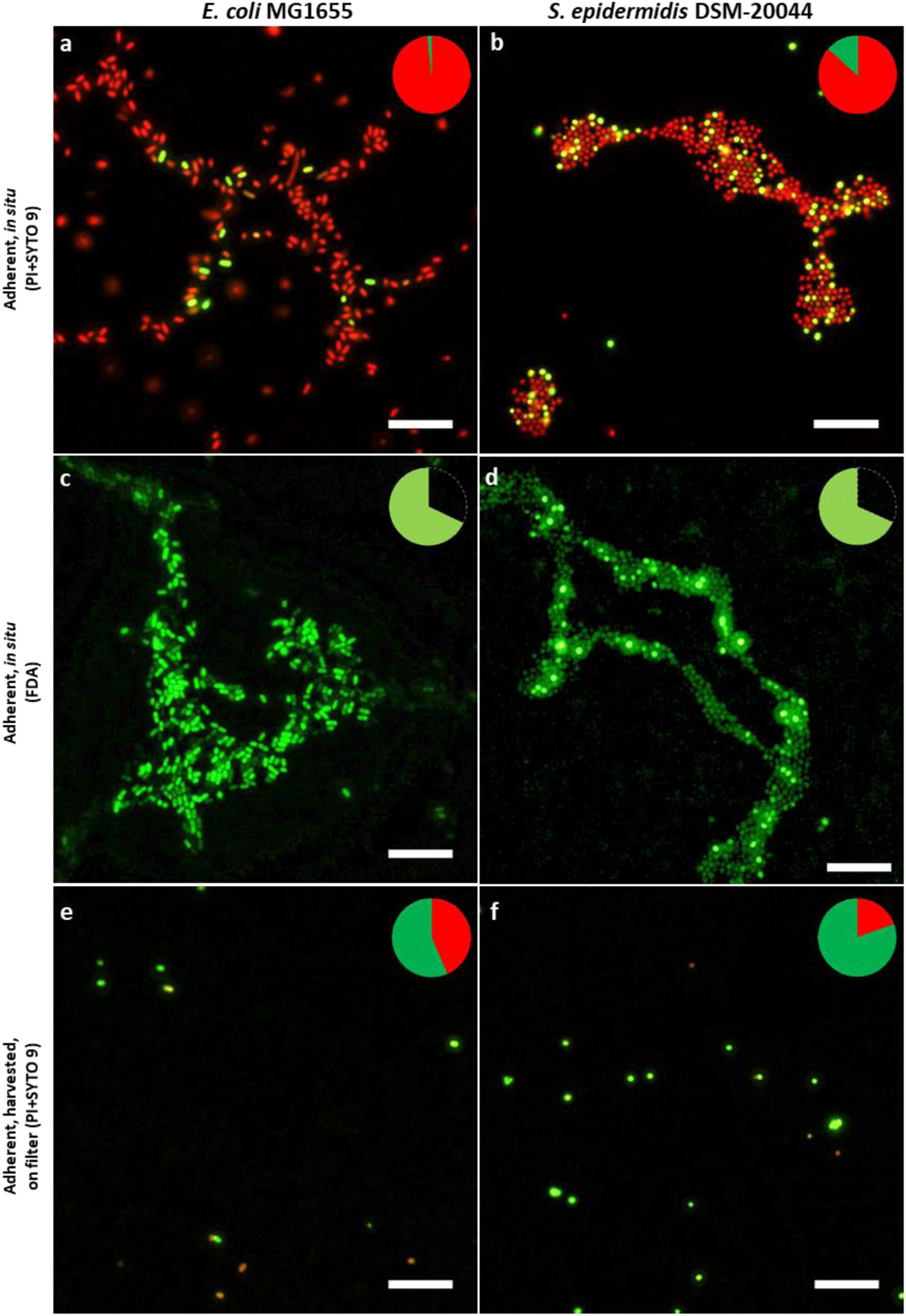
Epifluorescence microscopy images of adherent *E. coli* (a-d) and S*. epidermidis* (e-h) viability staining. 24 h initial monolayer biofilm formed on glass in PBS stained *in situ* with propidium iodide (PI) and SYTO 9 (a, b), with fluorescein diacetate (FDA) (c, d) or harvested via sonication, stained with PI and SYTO 9 and collected on filter (e, f). Pie diagrams represent total cell count on surfaces with PI, SYTO 9 and FDA stained signal proportions marked in red, dark green and light green respectively. Scale bars correspond to 10 µm.

**Figure 2.**
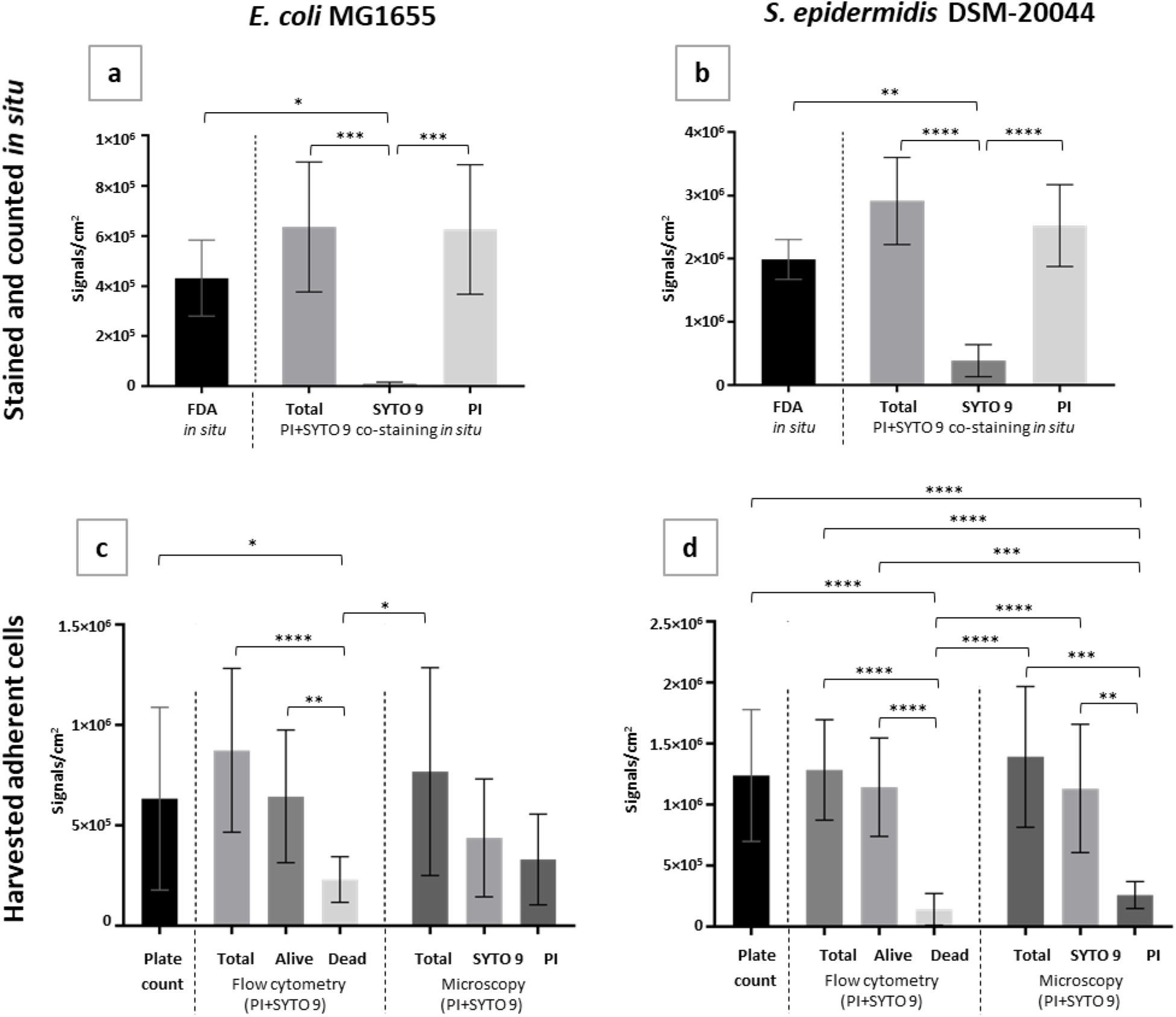
Comparison of multiple approaches to evaluate adherent cell viability in *E. coli* (a, c) and *S. epidermidis* (b, d) biofilms on surface *in situ* (a, b) or after harvesting via ultrasonication (c, d). 24 h initial monolayer biofilm formed on glass in PBS stained ***in situ* (a, b)** with propidium iodide (PI) and SYTO 9 or FDA followed by epifluorescence microscopy (EM) and signal counting or **harvested (c, d)** and cultivated for plate counts, co-stained with PI and SYTO 9 and analyzed by flow cytometry (FCM) or collected on filter followed by EM and signal counting. Results are presented as signals/cm^2^ where one signal counted corresponds to a single fluorescent cell or diplococcus (microscopy), a CFU (cultured) or a FCM event. Live/dead gating of FCM signal populations was based on known proportions of viable and ethanol-killed planktonic bacteria. Mean and standard deviation of 4-6 independent values for *in situ* staining and filtering and 10-16 independent values for plate counts and FCM are shown and only statistically significant differences (p<0.05) marked on graphs (“ “>0.05; *<0.05; **<0.01; ***<0.001; ****<0.0001)

To reveal the metabolic activity of *E. coli* and *S. epidermidis* in biofilms, we also stained the adherent cells with fluorescein diacetate (FDA), not a DNA-binding, but enzymatic activity indicative stain that emits green fluorescence after intracellular enzymatic cleavage ^37^. It was observed that 67.91% *E. coli* and 68.30% *S. epidermidis* cells were metabolically active compared to *in situ* total counts (Fig. 1c, 1d, 2a, 2b). Comparison of the results from staining the cells with FDA and PI+SYTO 9 showed that for both species of bacteria, there is a statistically significant difference in FDA and SYTO 9 signal counts (Fig. 2a, 2b) but there is no significant difference between FDA and PI or total (PI+SYTO 9) signal counts. On the assumption that dead cells are not metabolically active, and starvation may even cause underestimation of viable cell count based on FDA staining, this result sharply contradicts PI+SYTO 9 viability staining results. From these results demonstrating that most of the cells on glass surfaces are metabolically active and stain with PI while a minority of presumably viable cells stain with SYTO 9 only it can be concluded that SYTO 9 signal count significantly underestimates viability and PI signal count significantly overestimates dead cell count as a result of PI+SYTO 9 co-staining *in situ*. However, neither membrane integrity nor enzymatic activity, especially when incubated in a nutrient-poor environment, can truly indicate the reproduction capability of the cells. This can only be measured by cultivation-based methods.

### Viability staining and cultivability of harvested cells

Adherent bacteria were harvested from the surfaces via ultrasonication optimized to acquire the maximum number of viable cells (Supplementary Fig. 2) and plated or stained with PI and SYTO 9 and analyzed by flow cytometry (FCM) or collected on filter followed by epifluorescence microscopy (Fig. 1e, 1f, 2c, 2d). Of the 96.35±5.30% *in situ* PI-positive *E. coli* cells, only 43.50±5.30% were PI-positive after harvesting, subsequent staining and collection on filter, and only 27.76±9.61% of those cells could be assigned to the “dead” gate in FCM, based on ethanol-killed planktonic cells. Similarly, of the 75.69±18.44% *in situ* PI-positive *S. epidermidis* cells, only 19.56±8.93% were PI-positive after harvesting, staining and collection on filter, and only 11.07±10.70 those cells could be assigned to the “dead” gate in FCM. This result showing increased fraction of SYTO 9 stained cells after harvesting of adherent cells via ultrasonication compared with adherent cells *in situ* (Fig. 1a vs 1e; 1b vs 1f) was rather surprising. One would expect that sonication does not increase but rather decreases cellular viability due to physical damage as longer sonication durations resulted in decreased planktonic cell viability as well as decreased viable yield of adherent cells (Supplementary Fig. 2). However, due to the seemingly reversed red to green ratio after ultrasonication we hypothesized that sonic treatment affects the staining of viable cells with PI. One of the possible explanations for that was partial removal of eDNA containing extracellular matrix (ECM) from adherent cells. Indeed, sonication is a technique that is commonly used for ECM extraction ^38,39^. Removal of eDNA and false dead signals along with ECM was further confirmed by cultivating the harvested bacteria. Following the PI+SYTO 9 staining principle, plate counts could be expected to be smaller than the number of SYTO 9 signals from *in situ* staining due to possible cell aggregates forming only one colony but yielding several signals counted. On the contrary, compared to total signal counts from harvested and PI+SYTO 9 stained samples at least 82.43% of *E. coli* and 89.02% of *S. epidermidis* cells were cultivable and formed colonies on nutrient agar.

There was no statistically significant difference in plate counts of biofilm harvested cells, FCM total event counts FCM “alive” event counts, and SYTO 9 counts of harvested, PI+SYTO 9 stained and filtered samples for neither species, indicating that the majority of the harvested cells are truly viable (Fig. 2c, 2d) and the fact that they stained red with PI in *in situ* biofilms was indeed an artifact, most likely due to the presence of eDNA in the biofilm matrix.

Of the approaches used for harvested cell viability assessment, FCM proved to be a quicker and less elaborate method than filtering stained samples and counting fluorescent signals from microscopy images but gating harvested sample signals in FCM proved to be problematic. FCM alive and dead gates were based on viable and ethanol-killed planktonic cultures but unlike planktonic samples from the same test system, sonicated biofilm samples had much higher noise level and less defined and/or shifted “alive” signal populations (Supplementary Fig. 3) easily explained by partial ECM removal during harvesting and resulting double-staining of viable bacteria to various degrees in contrast to more strictly PI-defined “dead gate”.

The fact that viability estimate based on *in situ* PI-staining was significantly lower than the ones based on *in situ* FDA staining or harvested cell plate count (Table 1) suggested that eDNA could indeed play a major role in false “dead” PI-staining of biofilm bacteria *in situ*. To further confirm the hypothesis, confocal microscopy was used to better visualize the PI and SYTO 9 co-stained bacterial biofilms.

**Table 1.** Viability estimates (%)* of 24 h biofilms acquired with different methods.

| **Table 1** |  |  |  |  |  |
| --- | --- | --- | --- | --- | --- |
| **Viability estimates (%)\* of 24 h biofilms acquired with different methods.** |  |  |  |  |  |
| Species | *In situ*<br>PI+SYTO 9 | *In situ*<br>FDA vs PI+SYTO<br>9 total count | Harvested,<br>PI+SYTO 9,<br>on filter | Harvested,<br>PI+SYTO 9,<br>flow cytometry | Harvested,<br>plate count vs<br>PI+SYTO 9 total<br>count on filter |
| *E. coli* | 3.65±5.30 | 67.91 | 56.50±5.30 | 77.20±9.60 | 82.43 |
| *S. epidermidis* | 24.31±18.44 | 68.30 | 80.44±8.93 | 88.90±10.70 | 89.02 |

\* Mean and standard deviation of 4-6 independent values for *in situ* staining and filtering and 10-16 independent values for plate counts and FCM are shown. Percentages calculated as ratios of mean values are presented without standard deviations.

### Confocal laser scanning microscopy (CLSM) of PI+SYTO 9 stained biofilms

CLSM results revealed that PI and SYTO 9 signals generally colocalized (Fig. 3c, 3d, 4c, 4d) except for the most intensely red cells that lacked green signal and were presumably true dead signals. In order to have a closer look into the cells and attempting to spatially distinguish between red and green signals in adherent cells, vertical images of Z-stacks were taken from cross sections of *E. coli* and *S. epidermidis* monolayer biofilms (Fig. 3a, 4a). It must be noted that the result was seriously affected by vertical resolution limit of CLSM due to bacterial cell size, however was still able to bring light to the fact that most of the cells of both species have green interiors under red PI-stained exteriors. The same effect can be seen on single images from *S. epidermidis* CLSM Z-stacks (Fig. 3b; Supplementary Fig. 4) but is not so clearly distinguishable for *E. coli* (Fig. 4b; Supplementary Fig. 5). *S. epidermidis* CLSM results are obtained with the same staining conditions used in epifluorescence microscopy and FCM analysis. To visualize heterochromatic staining of bacterial cell interiors and exteriors for *E. coli*, PI concentration had to be decreased 20x. Higher PI concentrations could not be used in this case probably due to CLSM resolution limitations and *E. coli* cell proportions compared to *S. epidermidis* diplococci.

**Figure 3.**
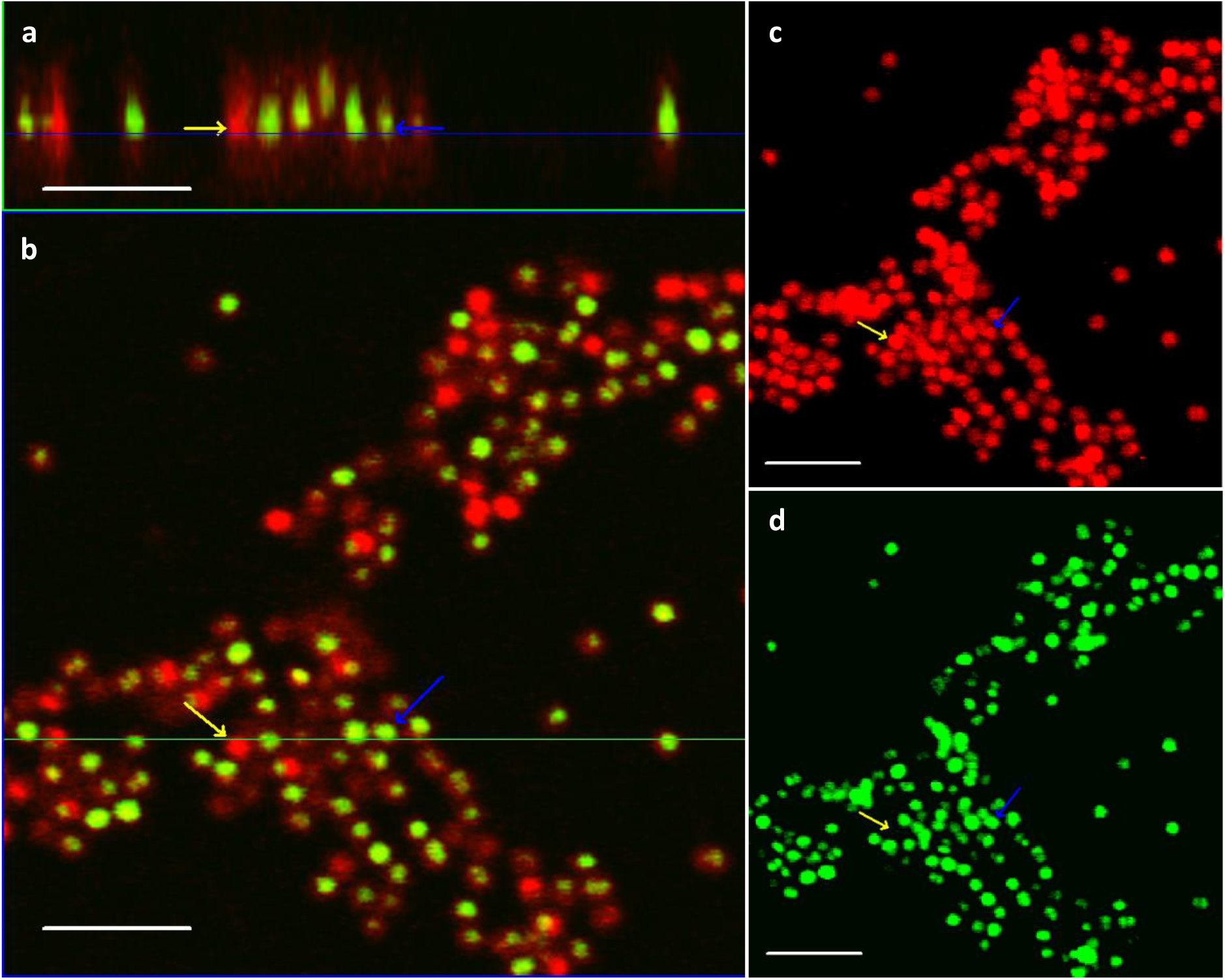
Confocal laser scanning microscopy (CSLM) images of 24 h *S. epidermidis* adherent cells in initial monolayer biofilm. co-stained with propidium iodide (PI) and SYTO 9: vertical cross-section from the Z-stack (a) and a single plane view from the same Z-stack (b); red channel (PI; c) and green channel (SYTO 9; d) maximum projections. Dead cells stained with PI are indicated with yellow and viable cells double-stained with PI and SYTO 9 with blue arrows. Scale bars correspond to 5 µm. Single images of the Z-stack are shown on Supplementary Fig. 4.

**Figure 4.**
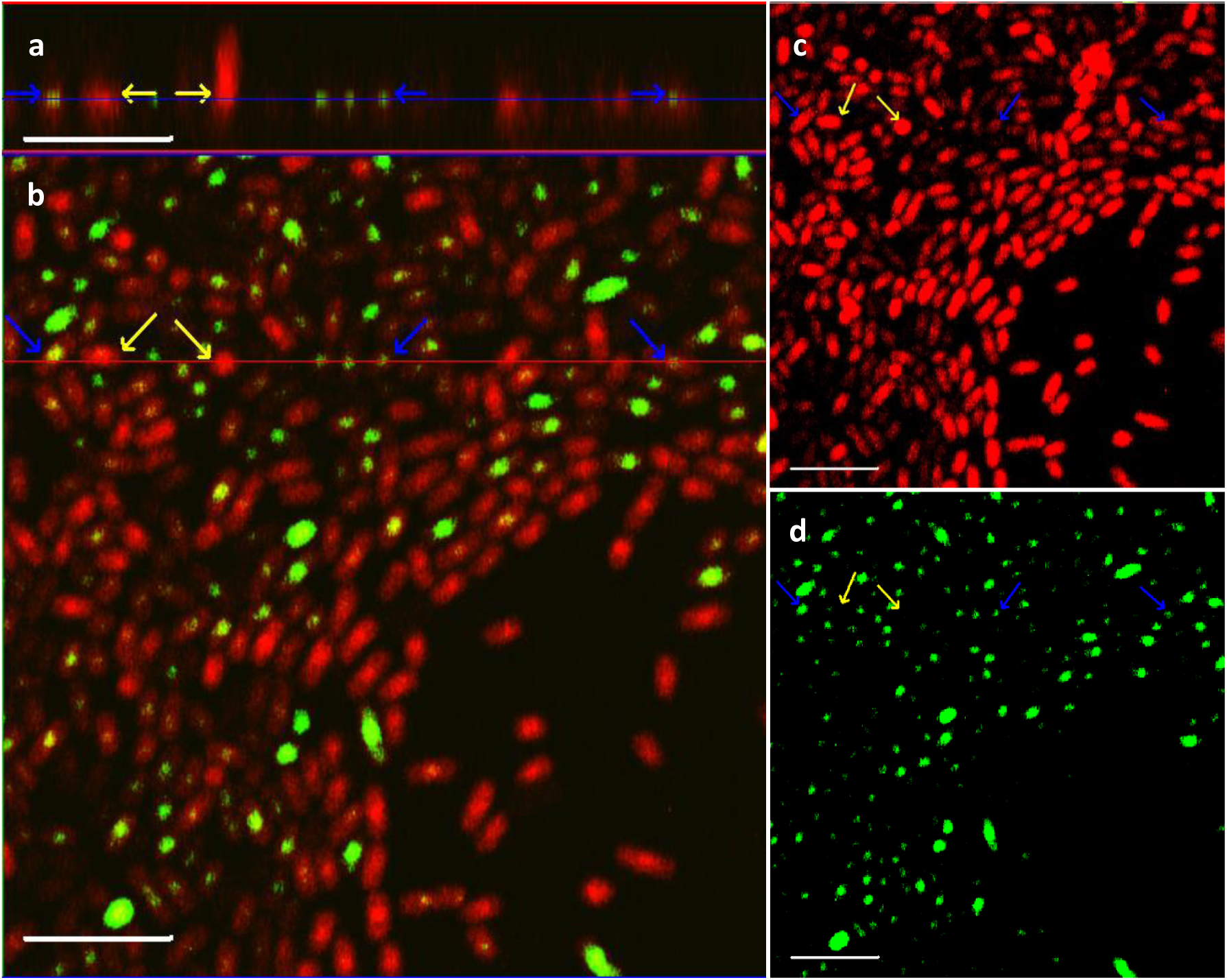
Confocal laser scanning microscopy (CLSM) images of 24 h *E. coli* adherent cells in initial biofilm. co-stained with PI and SYTO 9: vertical cross-section from the Z-stack (a) and a single plane view from the same Z-stack (b); red channel (PI; c) and green channel (SYTO 9; d) maximum projections. Dead cells stained with PI are indicated with yellow and viable cells double-stained with PI and SYTO 9 with blue arrows. Scale bars correspond to 5 µm. Single images of the Z-stack are shown on Supplementary Fig. 5.

As suspected from comparison of staining of cells with PI+SYTO 9 *in situ* and after sonication, CLSM confirms that PI really does stain cells externally and is therefore not indicative of membrane damage but produces false dead signals under these experimental conditions.

To further prove the role of eDNA in PI-staining, we also attempted to treat biofilms and similar amount of planktonic cells with DNase, remove the cells from the surfaces by scraping, concentrate by centrifugation and demonstrate larger amount of DNase degradable DNA signal on sessile cells than on planktonic cells. Unfortunately, the number of sessile cells optimized for *in situ* counting was too low to provide a signal in ethidium bromide agarose gel electrophoresis. Also, ECM removed from scraped cells by suspending them in 1.5 M sodium chloride ^40^ did not produce DNA signal on gel likely due to too low amount of DNA. PI+SYTO 9 staining and epifluorescence microscopy of 1.5M NaCl-treated cells confirmed that most of the cells indeed stained green suggesting successful removal of ECM, including eDNA, from the cells. It was also empirically observed that physical manipulations of adherent cells from scraping to centrifugation, vortexing and ultrasonication all shifted the red to green staining ratio to more green, indicating (partial) ECM removal in various steps of the process which makes cell number normalization between planktonic and low numbers of adherent cells prior to analysis difficult to achieve without losing significant amounts of eDNA.

## Discussion

The need to study possible false dead results of PI-based viability staining arose from our previous experiments carried out with bacterial biofilms in water and PBS, where we have similarly to this study, observed a large fraction of red PI-stained cells in biofilms on untreated glass but significantly smaller fraction of red-stained cells on glass surfaces with antibacterial treatment, although total cell counts on treated surfaces tended to be much lower than on untreated controls (unpublished data; Supplementary Fig. 6; surfaces described in ^41^). Yet the morphology of biofilms on untreated glass appeared normal while on antibacterial glass surfaces the biofilm structure as well as *S. epidermidis* characteristic diplococcal aggregation was disturbed. This result suggested almost reverse staining of alive and dead bacterial cells with PI and SYTO 9 in biofilms. In this study we show that in similar conditions on untreated glass membrane integrity based viability staining with DNA-binding propidium iodide can and will significantly overestimate the dead cell count in 24 h gram-positive and gram-negative monospecies biofilms in PBS. 96.35±5.30% of *E. coli* and 75.69±18.44% of *S.epidermidis* cells stained red and according to general viability staining principles could be considered “dead” when co-staining with PI and SYTO 9 *in situ* compared to 67.91% *E. coli* and 68.30% *S. epidermidis* being FDA-positive – metabolically active *in situ,* and at least 82.43% of *E. coli* and 89.02% of *S. epidermidis* cells being cultivable after harvesting from biofilms via ultrasonication. It was also evident that the red to green signal ratio was reversed after sonication which indicates that (partially) removed ECM during physical manipulation of the cells played a role in this process.

Compared to harvested cell plate count, FDA staining results *in situ* seem to underestimate viability (Table 1). This could be due to a few reasons. Firstly, the FDA method is challenging to work with due to weak fluorescent signals that require long exposures leading to photobleaching, high background fluorescence and varying signal intensities between individual cells (especially in the case of *S. epidermidis*). Secondly, FDA is indicative of metabolic activity, but biofilms were formed in a very nutrient-poor environment in which metabolic activity is expected to be slowed and as shown by Chavez de Paz et al. for oral bacteria, can reversibly affect FDA staining outcome ^42^.

Confocal laser scanning microscopy revealed double-stained cells with green fluorescing interiors under red stained exteriors of individual cells and confirmed PI staining not being indicative of membrane integrity but rather staining of eDNA which is one of the components of bacterial ECM. CLSM results look similar to what has been previously demonstrated, but not quantified, for *Bacillus cereus* biofilm on glass wool by Vilain et al ^43^, although in their study, the biofilm was formed in rich medium. Gallo et al ^44^ also noted a similar picture using surface amyloid fibre (SAF) producing and GFP-expressing *Salmonella typhimurium* biofilm cells surrounded by a “corona” of PI-stained eDNA and SAF complexes concluding that PI stains the cells externally. SAF and eDNA interactions have also been demonstrated for other species. For example, eDNA has been shown to facilitate the polymerization of SAF monomers in *Staphylococcus aureus* biofilms ^45^ and *E. coli* SAF monomer has been shown to bind to DNA, promoting SAF assembly ^46^. SAF and eDNA have been shown to facilitate bacterial attachment to surfaces and cell-cell aggregation ^47^. SAF and eDNA interactions in the context of biofilm formation and mechanical resistance need to be studied further to bring light to underlying mechanisms.

Moreover, the role of eDNA in PI-staining of adherent bacterial cells may not be constant in different biofilms but significantly affected by biofilm growth conditions. In our study we used biofilms grown in nutrient-poor PBS and no other conditions that could negatively affect adherent cultures were applied. However, in a more usual experimental setup, different treatments causing physical, toxic, starvation *etc*. stress to biofilms, especially in antimicrobial or anti-biofilm research together with a negative no-stress control are used. In the light of eDNA interfering with viability staining results, these stress factors could not only affect cell viability but also adherence efficiency and with that the amount of ECM and eDNA thereby potentially falsely exaggerating mortality of stress-treated samples compared to no-stress controls. For example, metabolic stress due to sub-lethal concentrations of antibiotics or other toxic compounds has been shown to enhance biofilm formation and/or result in higher eDNA content of the biofilms ^48–51^. Growth conditions, such as temperature, aerobic and starvation stress were reported to affect surface attachment and eDNA-mediated mechanism of biofilm formation of *Campylobacter jejuni* ^52^. DNase-sensitive eDNA dependent biofilm formation of *Streptococcus mutans* was observed in low pH stress and not in neutral pH ^53^. Higher eDNA content of biofilms subjected to physico-chemical stress was recently also observed for *S. epidermidis* ^54^. Not only severe stress, but also growth media selection can be of importance. Kadam et al. observed highest biofilm formation in nutrient-poor mediums and noticed that DNase-sensitive *Listeria monocytogenes* biofilms grown in nutrient broth consisted of clearly higher proportion of PI-positive cells during PI+SYTO 9 co-staining than DNase-insensitive biofilms of the same strains grown in a more nutrient rich brain heart infusion ^55^.

Together, this hints that the external staining phenomenon of PI might not only be dependent on the species used or starvation conditions but is also attachment-specific and dependent on conditions affecting matrix eDNA content, including different stress-responses.

## Conclusion

Viability estimation is of critical importance in evaluating antimicrobial/anti-biofilm surfaces and substances efficiency. Although the presence of extracellular nucleic acids in bacterial biofilm matrixes is well established in the literature, PI-based viability staining has remained a widely used tool for *in situ* viability estimate of adherent cells not taking into account possible eDNA interference in the viability staining results. From this study it can be concluded that membrane integrity based viability staining with DNA-binding dyes, including, but presumably not limited by PI, can significantly overestimate dead cell counts in the presence of eDNA in biofilms. To overcome this, the possible effect of eDNA should be controlled for by either: 1) using culture-based methods as a reference; 2) assess metabolic activity (e.g. esterase activity, respiration etc.) in parallel to DNA-staining and/or 3) minimizing extracellular matrix co-harvesting if harvested cell viability is to be assessed by staining. None of the aforementioned approaches are perfect for biofilms, but combination of methods rather than one approach is expected to result in more accurate estimations of viability.

## Methods

### Preparation of glass surfaces for bacterial attachment

18 mm x 18 mm soda-lime glass microscopy cover glasses (Corning, 2855-18) were used as biofilm carriers. Before inoculation carriers were rinsed with 70 vol% ethanol in deionized water and dried in biosafety cabinet with ultraviolet light irradiation for at least 20 minutes on both sides.

### Bacterial strains and biofilm cultivation

*S. epidermidis* DSM-20044 and *E. coli* MG1655 were grown overnight in Luria-Bertani broth (LB: 10 g/L tryptone, 5 g/L yeast extract in deionized water) at 30°C. Sterilized 18×18 mm glass cover slips were placed into wells of 6-well polycarbonate non-tissue culture coated plates (Corning, 351146). Bacterial cells where washed twice with PBS (180 mM sodium chloride, 3 mM potassium chloride, 9 mM dibasic sodium phosphate, 1,5 mM potassium hydrogen phosphate in deionized water, pH~7) using centrifugation at 7000 g for 10 min. Cell suspensions were immediately diluted to OD_600_ 0.01 in PBS and 5 ml of inoculum was pipetted onto glass surfaces in each well of the 6-well plates. Serial dilutions of remaining inoculum were made and drop-plated on nutrient agar (NA: 5 g/L meat extract, 10 g/L peptone, 5 g/L sodium chloride, 15 g/L agar powder in deionized water) to confirm inoculum cell count. Plates with inoculated surfaces were covered with lids and incubated at room temperature and ambient indoor lighting for 24 h to acquire biofilm density suitable for consecutive counting.

### Staining

Staining with PI (81845, Sigma) 20 mM and SYTO 9 (S-34854, Invitrogen^TM^ Thermo Fisher Scientific) 3.34 mM stock solutions in DMSO was carried out according to BacLight^TM^ Bacterial Viability Kit manual. Final concentrations of stains in 1:1 stain mixture in PBS was 30 µM PI and 5 µM SYTO 9 except for CLSM imaging of *E. coli* biofilm where PI concentration was reduced 20x to 1.5 µM. Stain mixture was either added to surfaces with biofilms (15 µl PBS-diluted stain mix pipetted straight onto surfaces and covered by cover slip), to cells harvested from surfaces by sonication or to planktonic bacteria collected from above the bacterial biofilms. The stained samples were incubated for 15 minutes in the dark (foil covered box) at room temperature.

FDA (201642, Sigma) stock solution used was 5 mg/ml in acetone, diluted 200-fold in PBS and kept on ice during the experiment. 15 µl of the stain solution was pipetted directly onto surfaces, covered by coverslip and incubated in the dark for 10 min before microscopy. Longer incubation periods yielded in higher background fluorescence and not significantly stronger signals from cells.

### Ultrasonication

Branson Digital Sonifier model 450 (max power 400 W) equipped with horn model 101-135-066R was used to harvest adherent cells from glass surfaces. The protocol was optimized to achieve maximal viable cell yield for sonication of glass surfaces in 50 ml glass beaker filled with 10 ml PBS at 25% sonication amplitude. For optimization, planktonic culture was used in parallel to biofilm and viability of both planktonic and harvested cells was evaluated during up to 30 sec sonication (Supplementary Fig. 2). Optimal time for sonication to achieve maximal viable cell yield was found to be 15 seconds for both bacterial species. Sonicated surfaces were stained with PI and SYTO 9 and microscoped to confirm removal of biofilm.

### Microscopy

Epifluorescence microscopy was carried out using Olympus CX41 microscope equipped with 100x oil immersion objective. Excitation filter cube DMB-2 (exciter filter BP475, dichroic mirror DM500, barrier filter O515IF) was used to filter mercury lamp emission allowing detection of both FDA as well as simultaneous detection of PI and SYTO 9 fluorescent signals. Images were captured with Olympus DP71 camera and Cell^B software, signal were counted in ImageJ software (“point” tool). For counting purposes at least 10 images were taken per sample at random locations.

Confocal laser scanning microscopy (CSLM) was carried out using Zeiss LSM 510 META equipped with 63x and 100x oil immersion objectives and acquired images analyzed in Zeiss LSM Image Browser. Blue 488 nm laser was used for excitation at 1.5% power and emission was observed in green channel (505-550 nm) and red channel (LP 575 nm). Z-stack heights imaged were 4.3 and 8.2 µm for *E. coli* and *S. epidermidis*, respectively, with 0.36 µm slice thickness.

### Flow cytometry

FCM analysis of PI and SYTO 9 co-stained bacteria was carried out using BD Accuri™ C6 device (BD Biosciences). Primary forward scatter (FCS-H) and secondary fluorescence signal (FL1-H) thresholds were used to filter out noise with minimal loss in bacterial cell signals and live-dead gating was done for *E. coli* and *S. epidermidis* using different proportions of viable overnight culture and ethanol-killed overnight culture (1h incubation in 70% ethanol) confirmed by plate counts. Gating of dead and alive signal populations was executed on SYTO 9 (FL1-A; 533/30 nm)/Propidium iodide (FL3-A; 670 nm LP) scatter plot as illustrated on Supplementary Fig 3.

### DNase treatment

15 surfaces per condition were prepared and rinsed as described and incubated with 500 µl 1x DNase I buffer (10x buffer: 100 mM Tris-HCl (pH 7.5), 25 mM MgCl_2_, 1 mM CaCl_2_) with or without DNase I (final concentration 100 U/ml, EN0523, Thermo Fisher Scientific). As a planktonic control, 3 ml of *E. coli* and 20 ml of *S. epidermidis* planktonic fraction, with estimated cell count similar to adherent cells on 15 surfaces were pelleted at 7000 g for 10 minutes, supernatant discarded and pellet suspended in DNase buffer with or without DNase I. Both, surfaces with biofilm and tubes with planktonic bacteria were incubated at 37°C for 4 hours. Adherent cells were harvested by scraping with cell scraper in the same buffer and pelleted by centrifugation at 7000 g for 10 min, suspended in 300 µl 1.5 M NaCl to remove extracellular matrix (ECM) as described in ^40^, thoroughly vortexed and pelleted again to remove cells from ECM fraction. 30 µl of ECM fraction in the supernatant was run on agarose gel electrophoresis (0.8% agarose in Tris-acetate-EDTA (TAE) buffer, stained with 0.5 μg/ml ethidium bromide; 60 V, 60 min, visualized on UV-transilluminator). Pelleted cells were resuspended in PI and SYTO 9 co-stain solution in final concentrations as described above and either analyzed by FCM or 5 µl pipetted onto microscopy slide, covered by cover slip, incubated in dark for 15 min and visualized with epifluorescence microscope.

### Statistical analysis

Mean values and standard deviations were calculated by Microsoft Excel standard functions. P-values used in Figure 2 were acquired using analysis of variance (ANOVA) followed by Tuckey’s multiple comparisons test at α=0.05 in GraphPadPrism 7.04 where analysis was executed individually for data presented on each graph (Fig. 2a-d).

## Supporting information

## Funding and acknowledgements

This work was supported by the Estonian Research Council grants IUT 23-5, PUT 748, European Regional Development Fund project TK134 and ERDF project Centre of Technologies and Investigations of Nanomaterials (NAMUR+, project number 2014–2020.4.01.16–0123) and EU COST Action CA15114 “Anti-Microbial Coating Innovations to prevent infectious diseases” (AMICI). NFA also acknowledges project POCI-01-0145-FEDER-006939 (Laboratory for Process Engineering, Environment, Biotechnology and Energy – UID/EQU/00511/2013), funded by the European Regional Development Fund (ERDF) through COMPETE2020 – Programa Operacional Competitividade e Internacionalização (POCI), and by national funds (PIDDAC) through FCT – Fundação para a Ciência e a Tecnologia/MCTES; project “LEPABE-2-ECO-INNOVATION” – NORTE–01–0145–FEDER–000005, funded by Norte Portugal Regional Operational Programme (NORTE 2020), under PORTUGAL 2020 Partnership Agreement, through the European Regional Development Fund (ERDF).

## Author Contributions

M. Rosenberg executed the experiments, participated in planning of the experiments and writing the paper. A. Ivask and N. F. Azevedo participated in planning of the experiments, writing the paper and providing background information on the subject.

## Competing Interests statement

Authors have no competing interests to declare.

## Data availability

The data generated in the current study is available from the corresponding author on reasonable request.

