## Supplementary material for "Propidium iodide staining underestimates viability of adherent bacterial cells"

### Supplementary figures

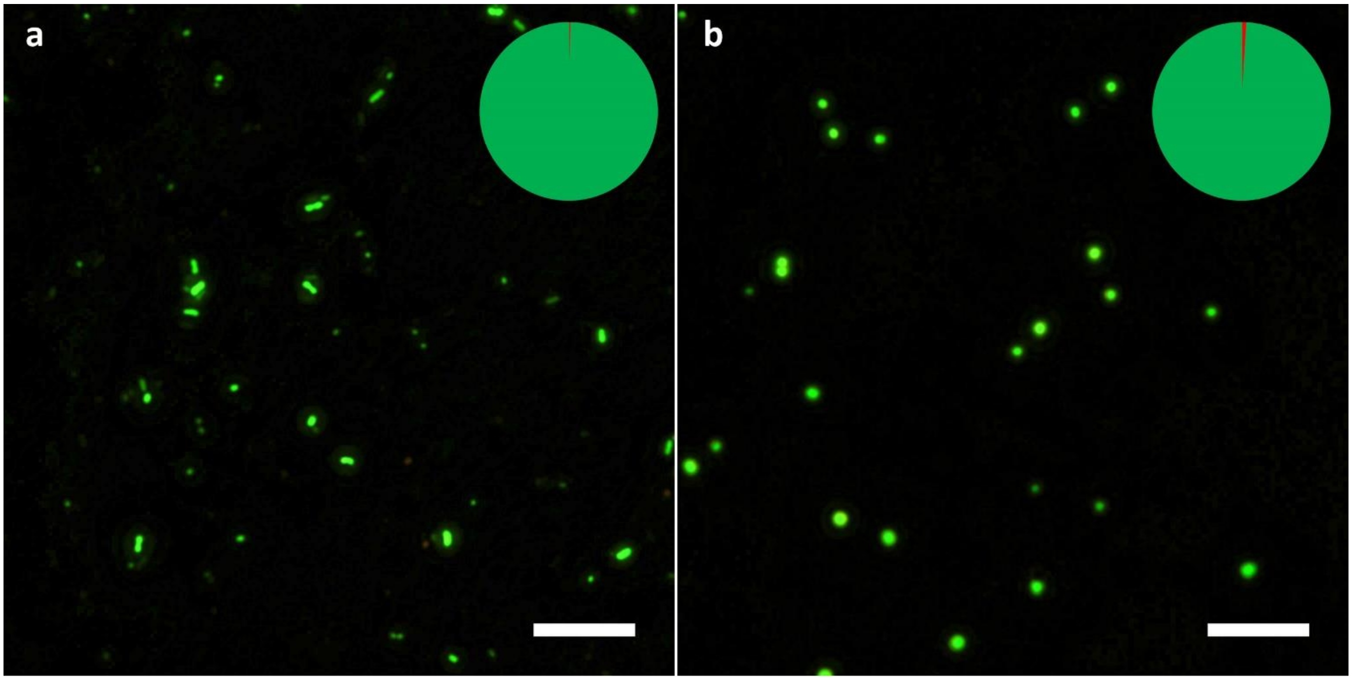


**Supplementary Figure 1**
**Epifluorescence microscopy images of planktonic *E. coli* (a) and S. *epidermidis* (b) viability staining:** planktonic cells from the biofilm experiment after 24 h incubation in phosphate buffered saline (PBS), stained with PI and SYTO 9 and collected on filter. Pie diagrams represent total cell count with PI, and SYTO 9 stained signal proportions marked in red and dark green respectively. Scale bars correspond to 10 µm.

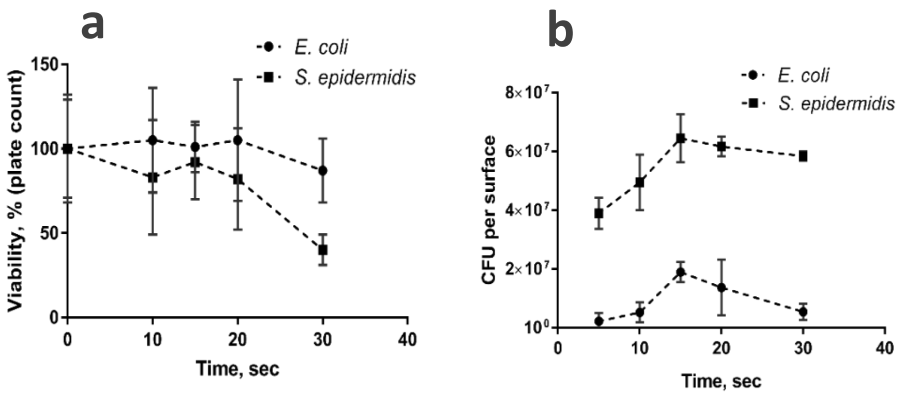


**Supplementary Figure 2**
**Optimization of protocol for ultrasonication of adherent bacteria from biofilms on glass surfaces**. Viability of planktonic bacteria ($OD_{600}$=0.05 in 10 ml PBS) during 0, 10, 15, 20 and 30 seconds sonication at 25% amplitude (a) and release of viable cells/aggregates from 24 h biofilm ($OD_{600}$=0.05 inoculum) in 10 ml PBS under the same conditions (b). Although planktonic culture endures longer treatment without decreasing viability, 15 seconds ultrasonication resulted in highest CFU per surface yield for biofilm cells. Glass surfaces were stained with propidium iodide and SYTO 9 after ultrasonication to confirm removal of biofilm.

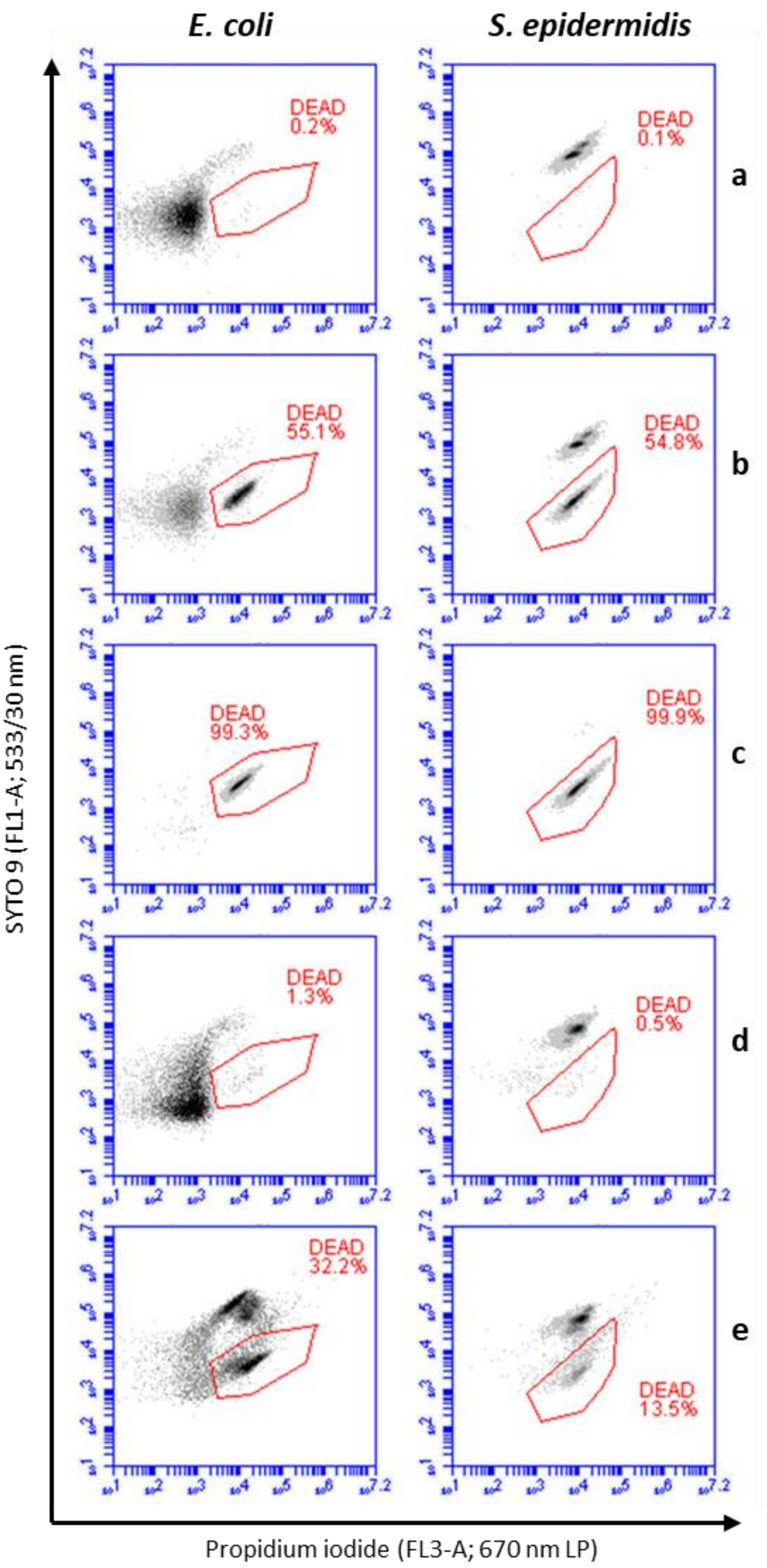


**Supplementary Figure 3**
**Flow cytometry (FCM) density plots of propidium iodide (PI) and SYTO 9 co-stained *E.coli* MG1655 and *S. epidermidis* DSM20044**: PBS washed overnight planktonic culture (a); PBS washed and ethanol-killed overnight culture (c); 1:1 mix of viable and ethanol-killed overnight cultures (b); pooled and ultrasonicated planktonic cells from biofilm experiment (d); ultrasonicated and pooled sessile cells from biofilm (e). Live/dead gating based on known proportions of viable and ethanol-killed planktonic bacteria was used to evaluate viability of bacteria harvested from 24 h biofilms. It is evident that gating strategy used is applicable for planktonic bacteria from the biofilm experiment even after incubating in PBS and sonication (d) while FCM populations of harvested *E. coli* biofilm cells have shifted and plots are much noisier. For adherent cells, dead gate seems to be better defined by PI staining than shifting populations of viable cells with presumably variable degrees of extracellular PI staining.

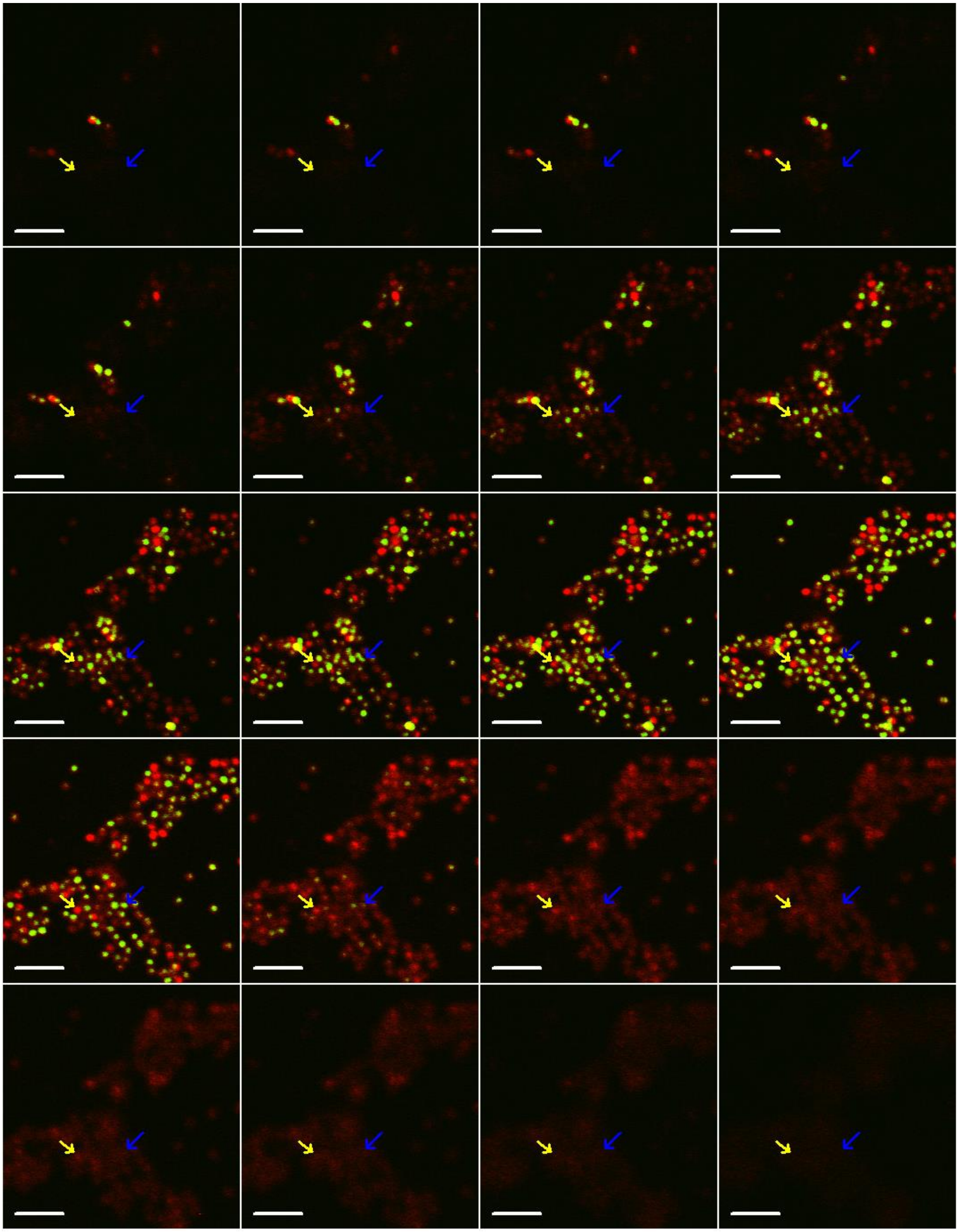

**Supplementary Figure 4**

**Confocal laser scanning microscopy (CLSM) stack images of 24 h *S. epidermidis* DSM20044 monolayer biofilm co-**stained with PI and SYTO 9. Dead cells stained with propidium iodide (PI) are indicated with yellow and alive cells double-stained with PI and SYTO 9 with blue arrows. Scale bars correspond to 5 µm.

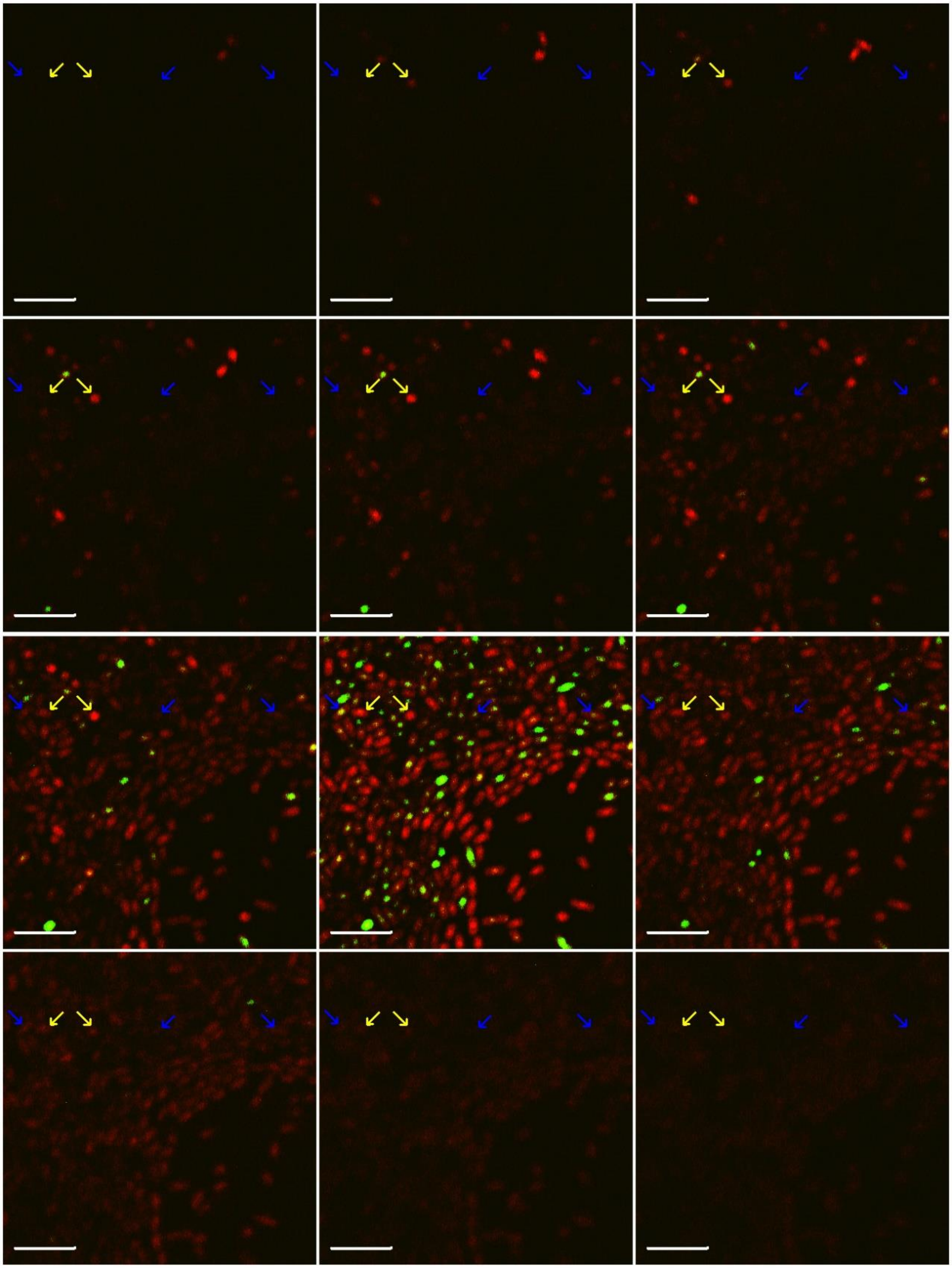

**Supplementary Figure 5**

**Confocal laser scanning microscopy (CLSM) stack images of 24 h *E. coli* MG1655 monolayer biofilm** stained with PI (concentration reduced to 1.5 µM) and SYTO 9. Dead cells stained with propidium iodide (PI) are indicated with yellow and alive cells double-stained with PI and SYTO 9 with blue arrows. Scale bars correspond to 5 µm.

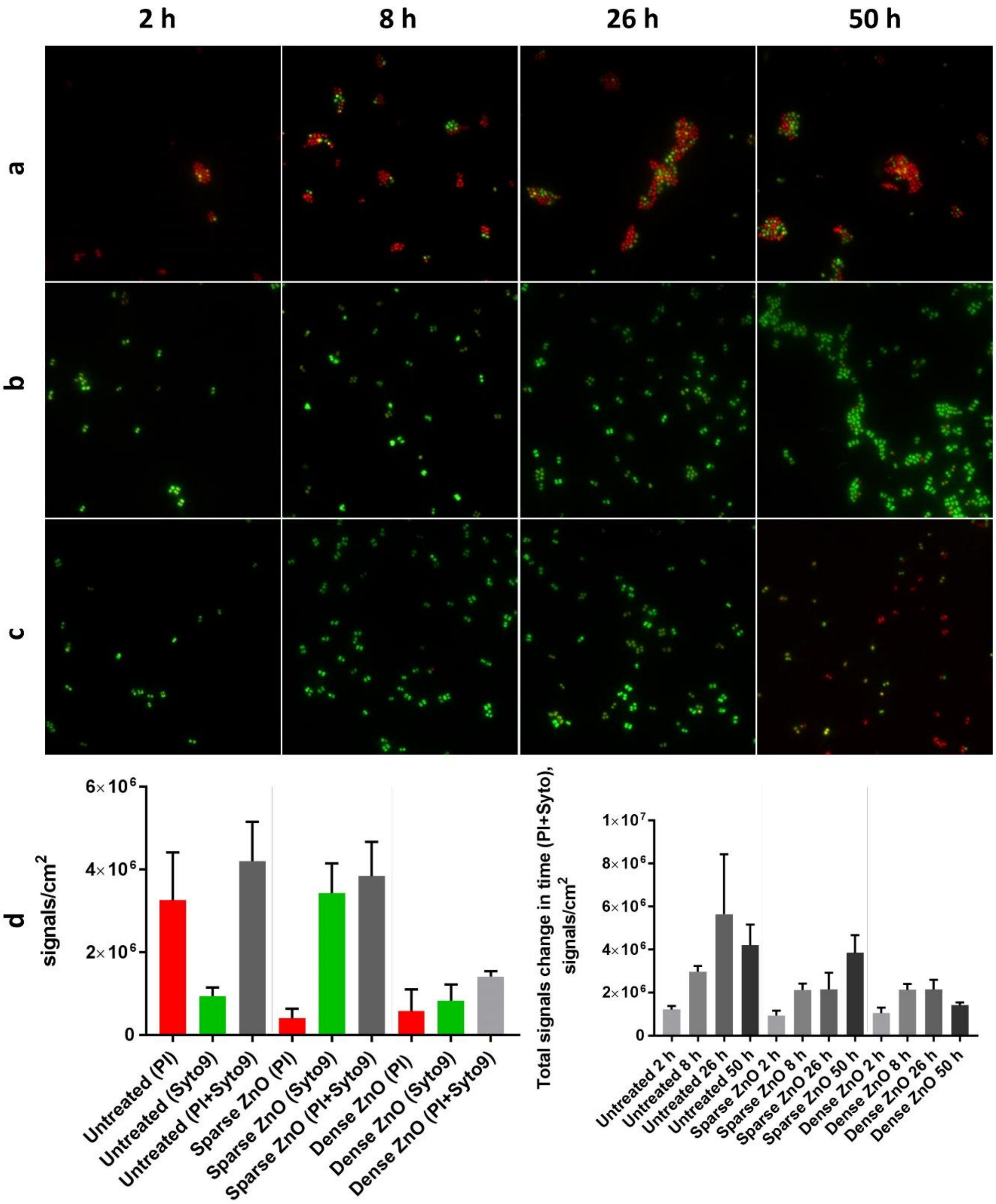


**Supplementary Figure 6**

**Staining of *S. epidermidis* biofilms on glass with propidium iodide (PI) and SYTO 9.** *S. epidermidis* DSM20044 biofilm on untreated glass (row a) and on antimicrobial nano-ZnO covered surface described in [51] (row b shows glasses with lower nano-ZnO content and row c shows glasses with higher ZnO content) in PBS stained *in situ* with PI and SYTO 9 and signal counts (row d) of 50 h biofilm (d, left) or total signal count change in time (d, right). Red staining is predominantly observed on non-toxic untreated glass with biofilm specific aggregates formed by tightly bound diplococci while on nano-ZnO antibacterial surfaces biofilm-specific aggregates appear later (b), but stain predominantly green and comprise of loosely bound diplococci and tetrads. Same staining pattern and forming loose diplococci and tetrads is true even in the case on surfaces with even higher nano-ZnO content where biofilm-specific aggregation is not observed during 50 h.
